# SGT1 is not required for bacterial PAMP-triggered immunity

**DOI:** 10.1101/2020.05.08.083766

**Authors:** Gang Yu, Liu Xian, Haiyan Zhuang, Alberto P. Macho

**Affiliations:** Shanghai Center for Plant Stress Biology, CAS Center for Excellence in Molecular Plant Sciences; Shanghai Institutes of Biological Sciences, Chinese Academy of Sciences, Shanghai 201602, China; University of Chinese Academy of Sciences, Beijing, China

## Abstract

Plant immune signaling activated by the perception of pathogen-associated molecular patterns (PAMPs) or effector proteins is mediated by PRRs and NLRs, respectively, and often share cellular components and downstream responses. The suppressor of the G2 allele of skp1 (SGT1) is a core immune regulator required for the activation of NLR-mediated immunity. In this work, we examined the requirement of SGT1 for PRR-mediated immune responses in both *Nicotiana benthamiana* and Arabidopsis. Using complementary genetic approaches, we found that SGT1 is not limiting for early PRR-dependent responses or anti-bacterial immunity. Therefore, we conclude that SGT1 does not play a significant role in bacterial PAMP-triggered immunity.

## Introduction

Cell surface-localized pattern recognition receptors (PRRs) and intracellular nucleotide-binding and leucine-rich repeat domain-containing receptors (NLRs) represent two tiers of the plant surveillance system against invading organisms (Jones and Dangl 2006). Both types of receptors perceive, directly or indirectly, invasion patterns from pathogens (Cook et al. 2015): PRRs mediate the perception of conserved elicitors (PAMPs; Pathogens-Associated Molecular Patterns), and their activation leads to the development of PAMP-triggered immunity (PTI) (Boller and Felix 2009); NLRs perceive effector proteins delivered inside plant cells, and their activation leads to effector-triggered immunity (ETI) (Jones and Dangl 2006; Lolle et al. 2020). Both PTI and ETI confer disease resistance. The activation of both PRRs and NLRs leads to convergent responses, such as ion fluxes, activation of MAP Kinase cascades, production of reactive oxygen species (ROS), as well as transcriptional reprograming, and therefore rely on similar or shared signaling components (Peng et al. 2018). Moreover, it has been recently reported that PTI and ETI require each other to confer robust disease resistance (Ngou et al. 2020; Yuan et al. 2020). The suppressor of the G2 allele of skp1 (SGT1), as a core immune regulator, is required for the activation of NLR-mediated immune responses (Shirasu 2009). In this work, we examined the requirement of SGT1 for PTI responses in both *Nicotiana benthamiana* and Arabidopsis. Although the Arabidopsis *SGT1* mutants showed different ROS burst after PAMP treatment, other PTI responses, including MAPK activation or anti-bacterial immunity, were not altered upon *SGT1* mutation or gene silencing. Therefore, we concluded that SGT1 does not play a significant role in bacterial PAMP-triggered immunity.

## Results and discussion

We recently found that SGT1s from different plant species are targeted by an effector protein injected into plant cells by the bacterial pathogen *Ralstonia solanacearum* (Yu et al. 2020). Such effector, named RipAC, interferes with the MAPK-mediated phosphorylation of SGT1 to suppress ETI responses (Yu et al. 2020). In an independent approach to identify additional activities of this effector, we found that RipAC is able to suppress the fast ROS burst triggered by bacterial PAMPs in *N. benthamiana*; RipAC expression almost completely blocked the early production of ROS upon treatment with the elicitors flg22^Pto^ (from *Pseudomonas syringae*; Figures 1A and 1B) or csp22^Rsol^ (from *R. solanacearum*; Figures 1C and 1D) (Wei et al. 2018). Considering that RipAC suppresses ETI by targeting SGT1 and our new findings that RipAC also suppresses PAMP-triggered responses, we considered a potential role of SGT1 in the activation of PTI. Although SGT1 is transcriptionally induced by different stimuli, including PAMP perception (Azevedo et al. 2006; Noël et al. 2007), whether SGT1 is required for PTI has not been formally tested to date. To address this question, we first performed virus-induced gene silencing (VIGS) of *NbSGT1* in *N. benthamiana* (Yu et al. 2019). Our *NbSGT1* VIGS approach results in undetectable levels of NbSGT1 protein (Figure S1A), and abolishes SGT1-dependent ETI responses, such as the cell death triggered by the *R. solanacearum* effector RipE1 (Sang et al. 2020; Figure S1B). However, VIGS of *NbSGT1* did not have a significant impact in the ROS burst (Figures 1E-1G) or MAPK activation (Figures 1H and 1I) triggered by flg22^Pto^ or csp22^Rsol^, indicating that SGT1 is not required for early responses triggered by bacterial PAMPs in *N. benthamiana*. Moreover, the ability of RipAC to suppress PAMP-triggered ROS was not affected in tissues that do not accumulate NbSGT1 (Figures 1J-1Q and S1C), confirming that RipAC suppresses PAMP-triggered ROS in an SGT1-independent manner.

**Figure 1.**
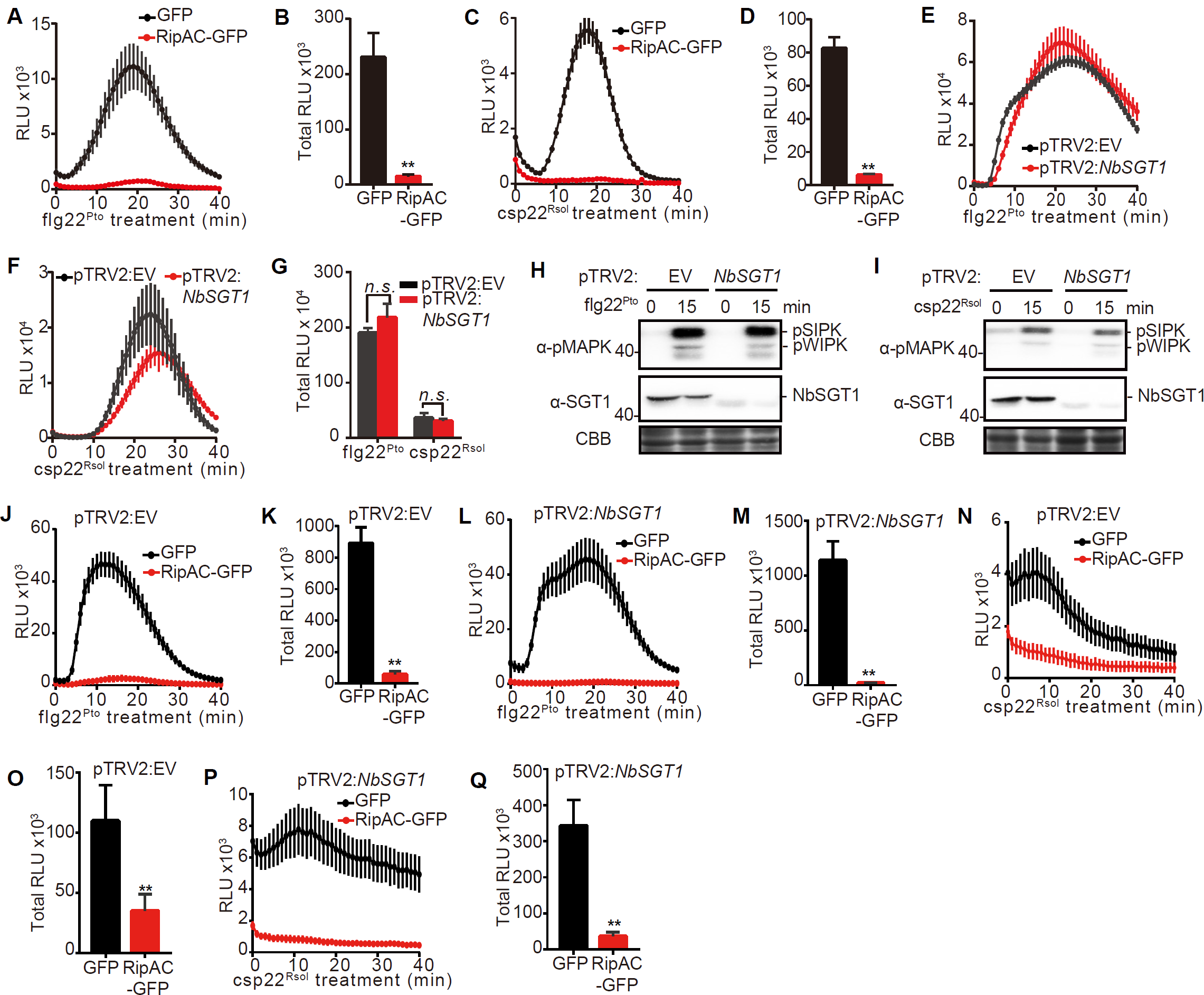
RipAC inhibits PAMP-triggered ROS in *Nicotiana benthamiana* leaves, which does not require NbSGT1. (A-D) RipAC suppresses ROS burst triggered by flg22^Pto^ (A and B) and csp22^Rsol^ (C and D). RipAC-GFP or GFP (as control) were transiently expressed in 5-week-old (for 50 nM flg22 ^Pto^) or 6-week-old (for 50 nM csp22^Rsol^) *N. benthamiana* leaves and ROS burst was analyzed after treatment with the respective elicitors using a luminol-based assay (mean ± SEM, n=24, *t*-test, **p<0.01). (E) and (F) Silencing *NbSGT1* does not affect PAMP-triggered ROS burst in *N. benthamiana*. PAMP (50 nM flg22^Pto^, or 50 nM csp22^Rsol^)-triggered ROS burst assay was performed 2-week-post *NbSGT1* VIGS using a luminol-based assay (mean ± SEM, n=16, *t*-test, *n.s*. indicates no statistical significance). Empty vector (pTRV2:EV) was used as control. (H) and (I) Silencing *NbSGT1* does not affect PAMP-triggered MAPK activation in *N. benthamiana*. PAMP (100 nM flg22^Pto^, or 1 uM csp22^Rsol^)-triggered MAPK activation was analyzed 2-week-post *NbSGT1* VIGS using an anti-pMAPK antibody. Empty vector (pTRV2:EV) was used as control. (J-Q) Silencing *NbSGT1* does not affect PAMP-triggered ROS production suppression activity of RipAC in *N. benthamiana*. RipAC suppresses ROS burst triggered by 50 nM flg22^Pto^ in either control plants (pTRV2:EV, J and K) or *NbSGT1* VIGS plants (L and M). RipAC also suppresses ROS burst triggered by 50 nM csp22^Rsol^ in either control plants (pTRV2:EV, N and O) or *NbSGT1* VIGS plants (P and Q). RipAC-GFP or GFP (as control) were transiently expressed in *N. benthamiana* leaves 2-week-post VIGS and ROS burst was analyzed after treatment with the respective elicitors using a luminol-based assay (mean ± SEM, n=8, Student *t*-test, **p<0.01). In A, C, E, F, J, L, N, and P, the graphs show ROS dynamics after PAMP treatment. In B, D, G, K, M, O, and Q, total RLU was calculated within 60 min after PAMP treatment. In (H) and (I), the western blots were probed with the antibodies indicated in the figures. The accumulation of endogenous NbSGT1 was detected using an anti-SGT1 antibody. Coomassie brilliant blue (CBB) staining was used as loading control. Molecular weight (kDa) marker bands are indicated for reference. All these experiments were performed at least 3 times with similar results.

RipAC associates with the two isoforms of Arabidopsis SGT1: AtSGT1a and AtSGT1b (Yu et al. 2020). In Arabidopsis, an *Atsgt1aAtsgt1b* double mutant is embryo lethal, suggesting that both genes have redundant functions in plant development besides their contribution to immunity (Azevedo et al. 2006). In short day growth conditions, we observed that the knockout mutants *sgt1a-1* (Ws-0 background) and *sgt1b-3* (La-*er* background) show different developmental phenotypes compared to their respective WT controls (Figure 2A): while *sgt1a-1* mutant plants are larger than WT plants, *sgt1b-3* were smaller than WT plants. To test PTI responses, since Ws-0 does not perceive flg22 (Gómez-Gómez et al. 1999) and the perception of csp22 is restricted to certain solanaceous plants (Wang et al. 2016), we treated plant tissues with the elicitor peptide elf18 (Kunze et al. 2004) from either *P. syringae* or *R. solanacearum* (Lacombe et al. 2010). Surprisingly, the *sgt1a-1* mutant showed attenuated ROS burst, while the *sgt1b-3* mutant showed enhanced ROS burst, compared to their respective WT controls (Figures 2B-2I). It is noteworthy that, although *AtSGT1a* is transcriptionally induced after different biotic stress treatments, *AtSGT1b* transcripts stay almost unaltered upon the same treatments (Azevedo et al. 2006; Noël et al. 2007). Moreover, considering that the PAMP-triggered ROS results were inversely proportional to the size of the plants (relative to their respective controls), it is possible that these opposite results are due to developmental effects caused by the respective mutations. Despite these differences, none of these mutants showed altered MAPK activation triggered by elf18^Pto^ (Figures 2J and 2K). Given the different effect of *sgt1a* and *sgt1b* mutations on PAMP-triggered ROS burst, we tested whether they are affected in resistance against a PTI-inducing strain. Figures 2L and 2M show that none of the mutants presented altered resistance upon inoculation with a non-pathogenic *Pto ΔhrcC* mutant, suggesting that, although AtSGT1a and AtSGT1b may have a different contribution to early PTI signaling, they are not required for the establishment of PTI. To overcome the observed partial redundancy of AtSGT1a and AtSGT1b, we performed VIGS of *AtSGT1a* in *sgt1b-3* mutant plants (Figure 2N). *AtSGT1a* VIGS reduced the accumulation of AtSGT1a (Figure 2O), sufficient to render silenced plants more susceptible to the ETI-inducing strain *Pto* AvrRpt2 (Figure 2P), suggesting that these plants have impaired SGT1 functions in the establishment of ETI. On the contrary, the silenced plants were not more susceptible to a *Pto ΔhrcC* mutant, or to *Pto* DC3000 (Figures 2Q and 2R), suggesting that abolishing SGT1 function in Arabidopsis does not have a major impact in the establishment of PTI. Altogether, our results indicate that the targeting of SGT1 does not account for the RipAC-mediated suppression of PTI, and SGT1 is not required for PTI against bacterial pathogens.

**Figure 2.**
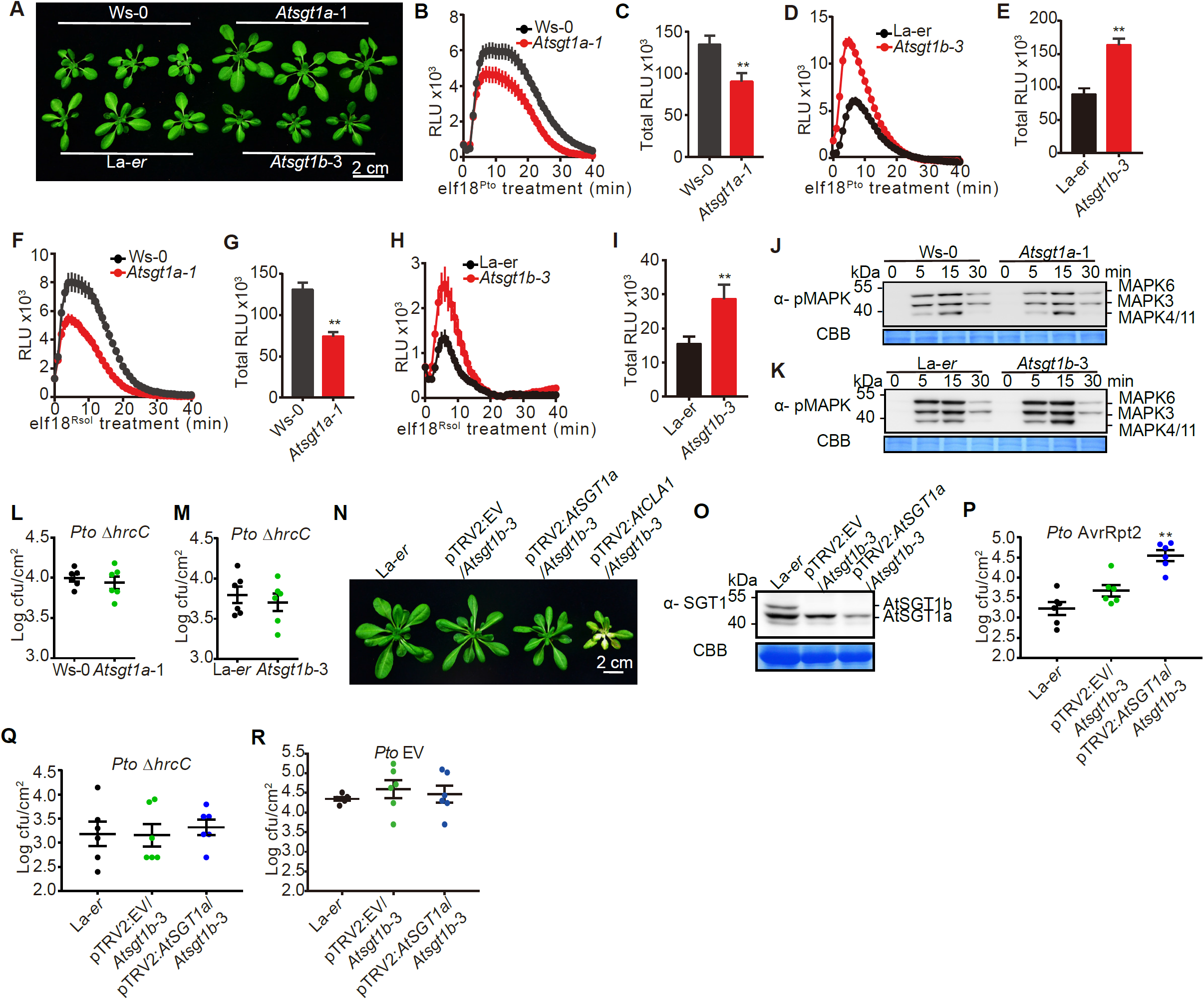
SGT1 is not required for PTI responses in Arabidopsis. (A) *AtSGT1a* and *AtSGT1b* mutants show different developmental phenotypes. The plants were grown in a short-day chamber (22°C, 10 h light/14 h dark photoperiod, 100-150 mE m^−2^ s^−1^, 65% humidity) and the pictures were taken 4-5 weeks after germination. (B-E) *AtSGT1a* or *AtSGT1b* mutants show different ROS burst after 100 nM elf18^Pto^ PAMP treatment. Leaf discs were taken from 4-5-week-old plants and the ROS burst was determined using a luminol-based assay (mean ± SEM, n=16, ** p<0.01, *t*-test). (F-I) *AtSGT1a* or *AtSGT1b* mutants show different ROS burst after 100 nM elf18^Rsol^ PAMP treatment. Leaf discs were taken from 4-5-week-old plants and the ROS burst was determined using a luminol-based assay (mean ± SEM, n=16, ** p<0.01, *t*-test). (J-K) Mutation of *SGT1* in Arabidopsis does not affect elf18^Pto^-triggered MAPK activation. 100 nM elf18^Pto^ was used to treat 12-day-old seedlings of the different genotypes and the samples were collected at the indicated time points. MAPK activation was analyzed using an anti-pMAPK antibody. (L-M) Mutation of *SGT1* in Arabidopsis does not affect *Pto ΔhrcC* bacterial growth. Arabidopsis plants of the different genotypes were hand-infiltrated with the non-pathogenic *Pto ΔhrcC* mutant strain, and four inoculated leaf discs were taken as one sample at 3 dpi (mean ± SEM, n=6). (N) Representative image showing *AtSGT1a* VIGS in *Atsgt1b*-3 mutant plants. The *AtCLA1* VIGS was used as a positive control, showing chlorosis caused by the silencing of *Arabidopsis Cloroplastos alterados 1* (AT4G15560). Two-week-old *Atsgt1b*-3 mutant plants were subjected to *AtSGT1a* VIGS and the pictures were taken 3-week-post VIGS. (O) Determination of *AtSGT1a* VIGS efficiency by western blot. Leaf samples were taken from different genotypes from (N) and protein accumulation was determined using an anti-SGT1 antibody. (P-R) Bacterial growth in *AtSGT1a* VIGS plants. Plants from different genotypes were hand-infiltrated with *Pto* AvrRpt2 (P), *Pto ΔhrcC* (Q), or *Pto* carrying an empty vector (EV, R), and four leaf discs from inoculated leaves were taken as one sample at 3 dpi (mean ± SEM, n=6, ** p<0.01, *t*-test). In B, D, F, and H, the graphs show ROS dynamics after PAMP treatment. In C, E, G, and I, total RLU was calculated within 60 min after PAMP treatment. These experiments were repeated at least 3 times with similar results. Coomassie brilliant blue (CBB) staining was used as loading control in (J, K, O).

Plants use diverse immune receptors to perceive invasion signals, leading to immune responses of similar nature, but different magnitude and duration (Peng et al. 2018). Although activation of both PTI and ETI leads to convergent downstream molecular events, the involvement and recruitment of different components before and after immune activation might be varied. In this work, we showed that, although extensive research has proven the requirement of AtSGT1b for the establishment of ETI (Azevedo et al. 2006; Shirasu 2009), SGT1 is not required for PTI responses in *N. benthamiana* or Arabidopsis. In this regard, SGT1 seems to be similar to the tomato NLR-required for cell death (NRC) proteins, which are NLR helpers required for ETI responses mediated by numerous NLRs, but are not essential for immunity triggered by bacterial flagellin (Wu et al. 2020).

## Author contributions

GY and APM designed experiments. GY, LX, and HZ performed experiments. GY and APM analyzed the data and wrote the manuscript.

## Acknowledgements

We thank Yasuhiro Kadota and Ken Shirasu for sharing biological materials and Xinyu Jian for technical and administrative assistance during this work. We thank the PSC Cell Biology core facility for assistance with confocal microscopy. This work was supported by the Strategic Priority Research Program of the Chinese Academy of Sciences (grant XDB27040204), the National Natural Science Foundation of China (NSFC; grant 31571973), the Chinese 1000 Talents Program, and the Shanghai Center for Plant Stress Biology (Chinese Academy of Sciences). Gang Yu was partially supported by the China Postdoctoral Science Foundation (grant 2016M600339). The authors have no conflict of interest to declare.

---

**Figure S1.**
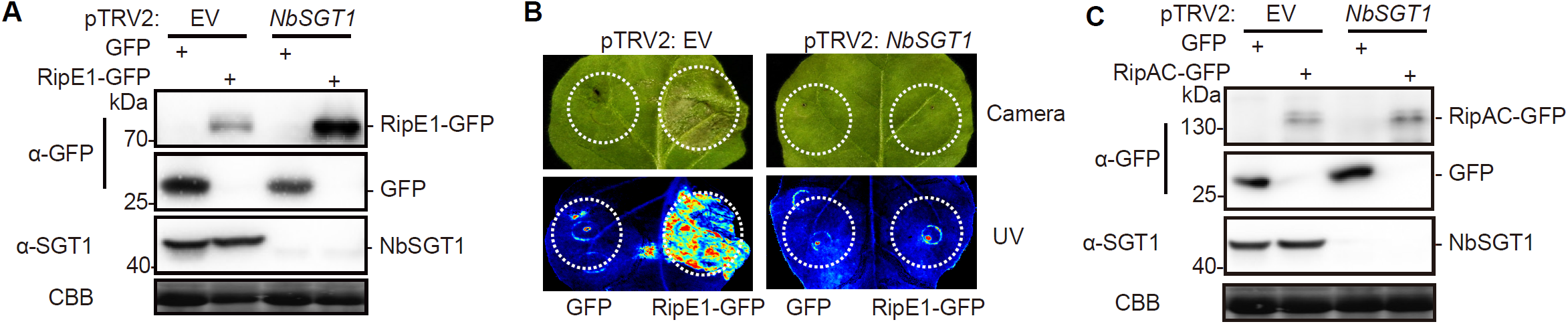
Silencing *NbSGT1* abolishes RipE1-triggered cell death in *N. benthamiana.* (A) Western blot showing the accumulation of GFP, RipE1-GFP, and endogenous NbSGT1 in (B). (B) Silencing *NbSGT1* abolishes RipE1-triggered cell death in *N. benthamiana.* RipE1-GFP or GFP (as control) were transiently expressed in the same leaf of *N. benthamiana* undergoing *NbSGT1* VIGS or VIGS with an empty vector (EV) construct (as control), using Agrobacterium with an OD600 of 0.5. Photos were taken 5 days-post-inoculation with a digital camera (upper panel) or a UV camera (lower panel). UV signal indicates the development of cell death (not GFP fluorescence). (C) Protein accumulation in Figure 1J-1Q. In (A) and (C), the western blots were probed with the antibodies indicated in the figures. The accumulation of endogenous NbSGT1 was detected using an anti-SGT1 antibody. Coomassie brilliant blue (CBB) staining was used as loading control. Molecular weight (kDa) marker bands are indicated for reference. All these experiments were performed at least 3 times with similar results.

## Supplemental Data

### Materials and Methods

#### Plant materials, constructs, and bacterial strains

The primers used to generate the constructs in this work are listed in the Supplemental Table S1. The *Atsgt1a*-1 (WS-0) (Azevedo et al. 2006) and *Atsgt1b-*3 (La-er) (Austin et al. 2002) have been described previously. In the experiments using Arabidopsis seedlings in 1/2 MS media, the seeds were grown on 1/2 MS plates in a growth chamber (22°C, 16 h light/8 h dark, 100-150 mE m^−2^ s^−1^) for germination and growth for 5 days, then the seedlings were transferred to 1/2 MS liquid media for another 7 days. For *Pseudomonas syringae* infection and reactive oxygen species (ROS) burst assays, Arabidopsis plants were cultivated in a short day chamber (22°C, 10 h light/14 h dark photoperiod, 100-150 mE m^−2^ s^−1^, 65% humidity) for 4-5 weeks. *Nicotiana benthamiana* plants were grown at 22°C in a walk-in chamber under 16 h light/8 h dark cycle and a light intensity of 100-150 mE m^−2^ s^−1^.

*P. syringae* pv. *tomato* (*Pto*) strains, including *Pto* containing an empty vector (EV), or a vector expressing AvrRpt2, or the type-III secretion system (T3SS)-defective mutant *Pto ΔhrcC*, were cultured overnight at 28 °C in LB medium containing 25 ug mL^−1^ rifampicin and 25 ug mL^−1^ kanamycin. *Agrobacterium tumefaciens* strain GV3101 with different constructs was grown at 28 °C on LB agar media with appropriate antibiotics. The concentration for each antibiotic is: 25 ug mL^−1^ rifampicin, 50 ug mL^−1^ gentamicin, 50 ug mL^−1^ kanamycin, 50 ug mL^−1^ spectinomycin.

#### Virus-induced gene silencing

Virus-induced gene silencing (VIGS) of *NbSGT1* in *N. benthamiana* was performed based on our previous report (Yu et al. 2019). Briefly, *A. tumefaciens* suspensions containing pTRV1 and pTRV2:EV or pTRV2:*NbSGT1* were mixed in 1 : 1 ratio to a final OD600=1 and infiltrated into two fully expanded leaves of 3-4-week-old plants (Liu et al. 2002). All the following experiments were performed 2 weeks after *NbSGT1* VIGS. The silencing efficiency was determined using western blot according to our previous report (Yu et al. 2019).

To silence *AtSGT1a* in *Atsgt1b-3* mutant background, VIGS was performed using pTRV vectors pTRV1 and pTRV2:*AtSGT1a* as previously described (de Oliveira et al. 2016), while the pTRV2:EV and pTRV2:*AtCLA1* were used as negative and positive controls, respectively. *A. tumefaciens* cultures were inoculated into LB medium containing 10 mM MES, 20 uM acetosyringone and appropriate antibiotics for overnight growth at 28°C. The cultures were collected and resuspended in the infiltration buffer with OD600=1.5 mixture (pTRV1:pTRV2-genes=1:1), which were then incubated at room temperature for 3 hours. The *A. tumefaciens* inocula were then inoculated into the first pair of true leaves of 2-week-old soil-grown *Atsgt1b-*3 mutant plants using a needleless syringe. The inoculated plants were covered overnight in a transparent dome in a growth room, and then transferred into a growth chamber for two to three weeks until the positive control, pTRV2-*AtCLA1*, showed plant albino phenotype. The silencing efficiency was determined with three random plants in each genotype by western using a custom anti-AtSGT1 antibody (Yu et al. 2019).

#### Transient gene expression, protein extraction, and immunoblot analysis

*A. tumefaciens* GV3101 carrying different constructs were infiltrated into leaves of 5-week-old wild-type (WT) or *NbSGT1* VIGS *N. benthamiana.* The OD600 used was 0.5 for each strain. To prepare the inoculum, *A. tumefaciens* was incubated in the infiltration buffer (10 mM MgCl_2_, 10 mM MES pH 5.6, and 150 uM acetosyringone) for 2 h.

To extract protein samples, 12-day-old Arabidopsis seedlings and leaf discs (diameter=18mm) from *N. benthamiana* were frozen in liquid nitrogen and ground with a Tissue Lyser (QIAGEN, Hilden, Nordrhein-Westfalen, Germany). Samples were subsequently homogenized in protein extraction buffer (100 mM Tris (pH 8), 150 mM NaCl, 10% Glycerol, 1% IGEPAL, 5 mM EDTA, 5 mM DTT, 1% Protease inhibitor cocktail, 2 mM PMSF, 10 mM sodium molybdate, 10 mM sodium fluoride, 2 mM sodium orthovanadate). The resulting protein samples were boiled at 70 °C for 10 minutes in Laemmli buffer and loaded in SDS-PAGE acrylamide gels for western blot. All the immunoblots were analyzed using appropriate antibodies as indicated in the figures. Molecular weight (kDa) marker bands are indicated for reference.

#### Pathogen inoculation assays, ROS burst assays, and cell death observations

For *Pto* inoculation, different *Pto* strains were resuspended in water at 1×10^5^ CFU mL^−1^. The bacterial suspensions were then infiltrated into 4-5-week-old Arabidopsis leaves. Bacterial numbers were determined 3 dpi.

ROS burst assays were performed in *N. benthamiana* and Arabidopsis plants as described previously (Sang and Macho 2017). To test whether RipAC suppresses PAMP-triggered ROS burst in *NbSGT1* VIGS *N. benthamiana*, the *A. tumefaciens* GV3101 carrying different constructs were infiltrated into leaves of *NbSGT1* VIGS or control (pTRV2:EV) *N. benthamiana* two days prior to leaf discs sampling. Plant leaf discs were collected and floated on sterile water overnight in 96-well plate. On the following day, ROS were elicited with 50 nM flg22^Pto^, 50 nM csp22^Rsol^, 100 nM elf18^Pto^ or 100 nM elf18^Rsol^, and the luminescence was measured over 60 min using a Microplate luminescence reader (Varioskan flash, Thermo Scientific, USA). Flg22^Pto^, csp22^Rsol^, elf18^Pto^, and elf18^Rsol^ peptide were purchased from Abclonal (China) and the peptide sequences are: flg22^Pto^: TRLSSGLKINSAKDDAAGLQIA; csp22^Rsol^: ATGTVKWFNETKGFGFITPDGG (Wei et al. 2018); elf18^Pto^: MAKEKFDRSLPHVNVGTI (Lacombe et al. 2010); elf18^Rsol^: MAKEKFERTKPHVNVGTI (Lacombe et al. 2010).

To observe the RipE1-triggered cell death, *A. tumefaciens* GV3101 carrying RipE1-GFP or GFP constructs were infiltrated into leaves of *NbSGT1* VIGS or the control (pTRV2:EV) *N. benthamiana* plants (Sang et al. 2020). The cell death phenotypes were recorded either by digital camera or a BIO-RAD GelDoc XR^+^ with Image Lab software (UV light) 5-d-post-inoculation (Yu et al., 2020). The leaf tissues were taken for western blots 2-d-post-inoculation.

#### MAP kinase activation assays

To evaluate the elf18^Pto^-triggered MAPK activation, two 12-d-old Arabidopsis seedlings were treated with 100 nM elf18^Pto^ and samples were collected at different time points. To evaluate PAMP-triggered MAPK activation in *NbSGT1* VIGS *N. benthamiana*, the plant leaves were treated with either 100 nM flg22 ^Pto^ or 1 uM csp22^Rsol^ for 15 min based on previous reports (Sang et al. 2018; Wei et al. 2018).

After protein extraction, the protein samples were separated in 10% SDS-PAGE gels and the western blots were probed with anti pMAPK antibodies (Phospho-p44/42 MAPK (Erk1/2), Cell Signaling, Cat# 4370) to determine MAPK activation. Blots were stained with Coomassie Brilliant Blue (CBB) to verify equal loading.

#### Quantification and statistical analysis

Statistical analyses were performed with Prism 7 software (GraphPad). The data are presented as mean ± SEM. The statistical analysis methods are described in the figure legends.

**Supplemental Table S1.**
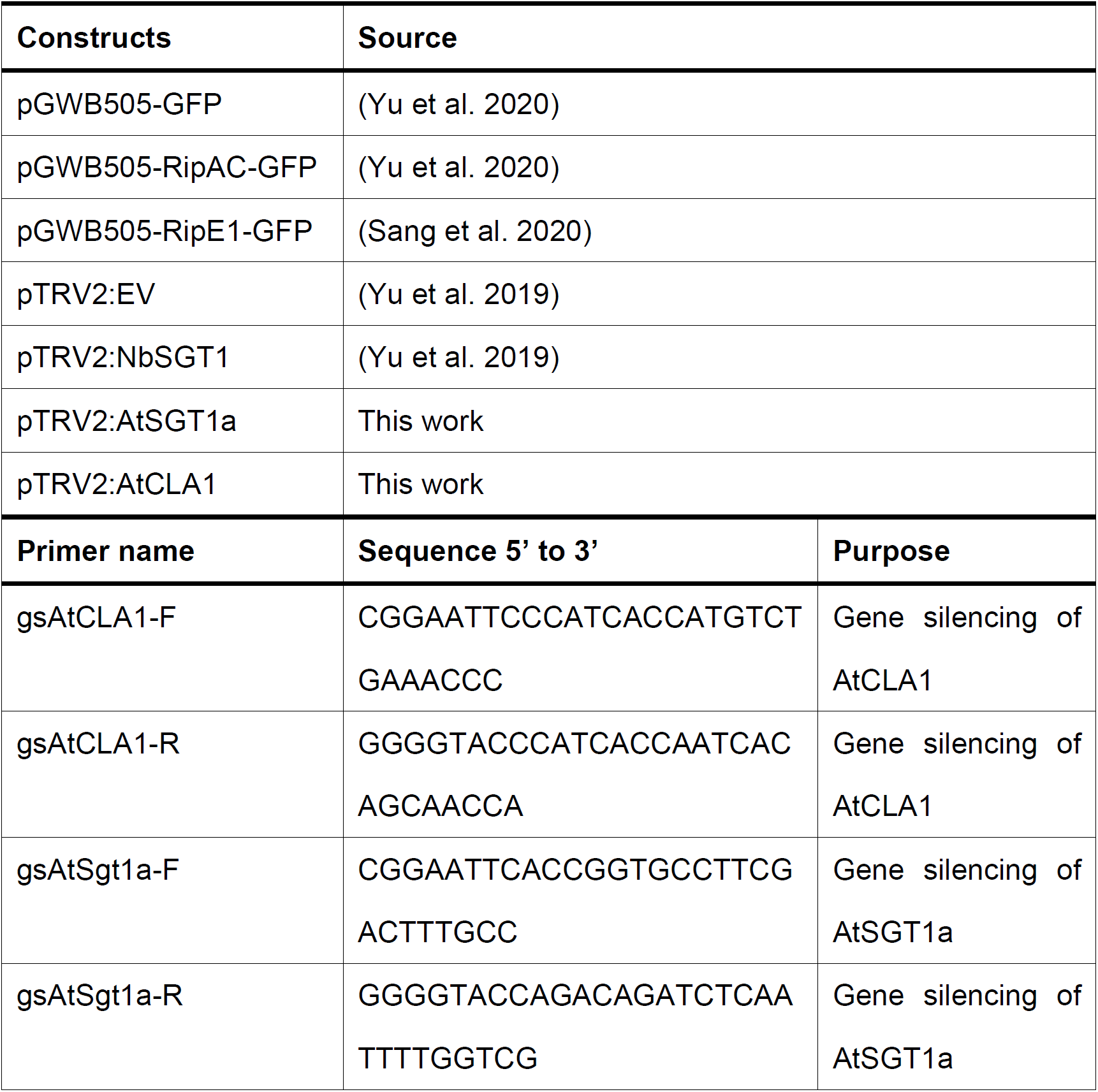
Constructs and primers used in this study

